# XPCLRS: fast selection signature detection using cross-population composite likelihood ratio

**DOI:** 10.64898/2026.02.27.708459

**Authors:** A. Talenti

## Abstract

**Summary:** The growing size of genomic datasets poses serious computational challenges, especially for laboratories and groups with limited access to high-performance compute facilities. This problem affects a broad range of analysis, including selection signature methods used to identify genomic regions undergoing selective pressure. Many of these methods were developed with SNP arrays in mind and are often not designed with scalability as a priority. Cross-population composite likelihood ratio (XP-CLR) is one such approaches, intended to detect hard selective sweeps by comparing two groups of individuals.

In this paper I introduce XPCLRS, a rust implementation of the XP-CLR selection signatures method. It can be hundreds of times faster, supports multithreading natively and produces results comparable to its original counterpart (R = 0.976). lowers the computational barrier to applying XP-CLR alongside other statistics, helping detect regions of interest, reduce false positives, and ultimately improve the robustness of genomic studies.

**Availability and implementation:** The source code for the software freely accessible in GitHub (https://www.github.com/RenzoTale88/xpclrs). Additionally, the software is also distributed via crates.io (https://crates.io/crates/xpclrs) and docker container (https://hub.docker.com/r/tale88/xpclrs) for ease of installation. The software is released under MIT open source license.

## Text

Selection signatures are a cornerstone of modern genomic analyses. They enable the detection of variants, genes and genomic regions that are under natural or human-driven selection, thereby revealing the genetic basis of key traits (de Simoni Gouveia *et al*. 2014). For example, selection signature analyses have proven valuable for identifying genes under selection related to productive traits (Ghildiyal *et al*. 2023; Deng *et al*. 2025), morphological characteristics (Plassais *et al*. 2017, 2019; Talenti *et al*. 2018), adaptation to environment (Tiwari *et al*. 2024) and convergent selection (Dutta *et al*. 2020) across different species.

Numerous methods have been developed to detect selection signatures, many of which focus on modelling selective sweeps – the process in which a beneficial allele rises in frequency and, through linkage disequilibrium, lead to the increase in frequency not of nearby neutral variants (Stephan 2019). These methods rely on different principles, and can be broadly grouped into within-population approaches (e.g. iHS; Voight et al. 2006), and comparative (cross-population) approaches (e.g. Fst; Holsinger & Weir 2009). Among comparative methods, the cross-population composite likelihood score (XP-CLR) method detects selective sweep by comparing allele frequencies between two populations to identifying signatures of selection. It models the neutral genetic drift as Brownian motion and uses it to identify sites that are diverging from it (Chen, Patterson and Reich 2010).

The method was first implemented in XPCLR, an open source Python software. It relies on multiple dependencies like numpy, scipy and scikit-allel to achieve faster performance (Harding 2025). The package performs well on low-to-medium density variant panels, but slows substantially when analysing larger datasets, an increasingly common occurrence. This is largely due Python being a high-level programming language: it enables rapid development and readable code, but imposes performance overheads compared with lower-level languages.

Rust is gaining traction among low-level programming languages thanks to its characteristics: near-C run times, a compiler that catches bugs at compile time, a built-in formatter improving readability of the code, the Cargo/crates ecosystem that simplifies dependencies management and software distribution, built-in support for multithreading, and memory safety that helps preventing security flaws. Bioinformatics has also rapidly adopted Rust, with tools appearing regularly – for example the rust-mdbg (Ekim, Berger and Chikhi 2021) and peregrine-2021 (Chin and Khalak 2019; Chin 2025) whole genome assemblers, Gfaffix to reduce graph genome complexity (Doerr and Marijon 2025), fastix to rename sequences in fasta files (Garrison 2025) and modkit to call and analyse modified bases from long read sequencing (Rand, Stoiber and Wright 2025).

Here, I introduce XPCLRS, a high-performance implementation of the XP-CLR method. Written in Rust, it uses the rayon crate for efficient multithreading (*Rayon - Crates*.*Io: Rust Package Registry* 2025) and scirs2-integrate for the quadrature integration (*Scirs2-Integrate - Crates*.*Io: Rust Package Registry* 2026). XPCLRS supports two formats: VCF, and its binary version BCF (Danecek *et al*. 2011), and binary PLINK (BED/BIM/FAM) (Purcell *et al*. 2007; Chang *et al*. 2015; Purcell and Chang), widely used to encode genotyping data generated from SNP array and sequencing studies alike. The tool provides the same settings of XPCLR, and expand by providing a `--fast` mode, where the quadrature integration is set to non-adaptive, providing further speed boost to the software.

I evaluated the performance and agreement of the tool with the original implementation. Using phased variants from the 1000 Genome project (https://ftp.1000genomes.ebi.ac.uk/vol1/ftp/release/20130502/), I ran XPCLR and XPCLRS on all biallelic sites extracted with bcftools (v1.23; options -m 2 -M 2) (Danecek *et al*. 2021; *Bcftools*) and analysed the two largest groups, European (EUR) and African (AFR). Figure 1A compares the results across modes – with and without the `--phased` and/or the `--fast` flags. To eliminate the randomness from marker subsampling and ensure a fair comparison, I set `--maxsnps` to 2000 to both pieces of software, so all markers in each window are used to estimate the statistics. The results are also collected in Supplementary Table 1 for manual inspection. Across all configurations, correlations were high (Pearson’s R = 0.955-0.976), confirming that XPCLRS reproduces XPCLR’s output. The small discrepancies stem from the numerical integration backends: XPCLR relies on scipy’s `quad` function calling QUADPACK’s adaptive general quadrature with singularities (QAGS), whereas XPCLRS uses the adaptive `quad` function from the scirs2-integrate crate.

**Figure 1.**
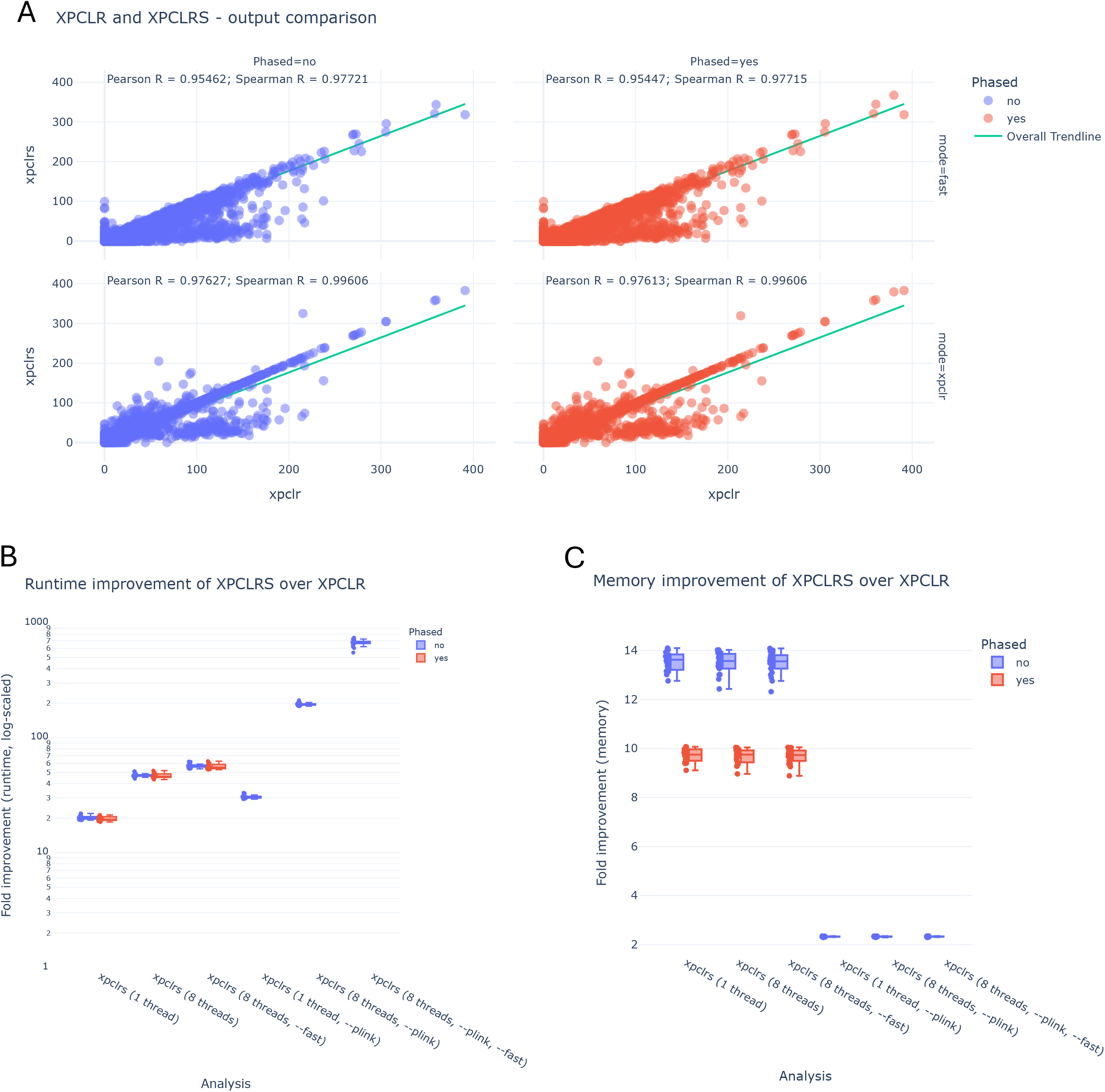
Comparison XPCLR and XPCLRS performance: A) correlations of the results generated by XPCLR and XPCLRS for phased/unphased and with/without the --fast option; B) log-scaled fold improvement in speed of XPCLRS over XPCLR across the different configurations for each chromosome; and C) fold improvement in memory usage of XPCLRS over XPCLR across the different configurations for each chromosome

While XPCLRS produces some differences in the values compare to XPCLR, these do not necessarily lead to a different interpretation of the results. Supplementary Figure 1 show the Manhattan plots for the absolute normalized XPCLR (pane a) and XPCLRS scores after genome-wide re-normalization 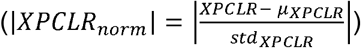 without and with `--fast` enabled (panes b and c, respectively). Results from XPCLR and XPCLRS show substantial equivalence in their peak distributions, unless `--fast` mode is enabled. This mode introduces greater differences in the output, particularly for weaker signals, making it more appropriate for preliminary analyses aimed at detecting only the strongest selection peaks.

Next, I compared the performance of XPCLRS with its original implementation, measuring wallclock time in seconds and peak memory usage using the GNU time utility in Linux. Runtime and memory results are shown in Figure 1B. The results of the same analysis is also reported in Supplementary Table 2. Figure 1B clearly shows the speed improvement achieved by XPCLRS, with a median of ∼19.4X improvement in the default single core model, and over 55X faster with 8-threads and `--fast` enabled. Some of the speed are masked by input/output (I/O) operations, which is similar for both XPCLR and XPCLRS using VCF as input format. However, this limitation is overcome by using PLINK as input format, further widening the speed gap to over 200X faster runtimes while still achieving high correlation (R=0.977, Supplementary Figure 2). Enabling `--fast` further increases throughput to 700X faster runtime, with a similarly modest drop in concordance (R = 0.955, Supplementary Figure 2). The memory usage of both tools are modest. Even so, Figure 1C shows the efficiency gains achieved by XPCLRS, using 8-14X less memory than XPCLR for the same dataset of 1,161 individuals, using well below 5GB peak RAM. Using the PLINK drops the gains to approximately 2x, effectively halving the memory requirement.

In addition to the runtime gains, XPCLRS differs from XPCLR in the way it handles multiallelic loci. XPCLR only considers sites with exactly two alleles globally – a reference and a single alternative allele. XPCLRS instead includes all sites that is biallelic within the sample analysed, even when both segregating alleles are non-reference. This expands the number of informative sites, particularly in diverse cohorts.

Beyond performance and feature differences, XPCLRS follows high coding standards. The GitHub repository uses continuous integration and development (CI/CD) to build and test across major target platforms (macOS and Linux, both x86 and ARM64 architectures), including end-to-end tests of the analyses. Each release automatically releases updated docker container and crates in crates.io. These distributions will simplify the adoption of the package and from the scientific community.

Finally, although the software is stable and fully usable, there it still room for improvement. Upcoming releases will reduce the memory footprint of reading PLINK files, add support other formats e.g. the newer PLINK2 binary format.

In conclusion, I present XPCLRS, a high-performance implementation of the XP-CLR method. XPCLRS expands the toolkit available to researchers by lowering the computational barriers and improving usability, assisting the discovery of novel candidate genes and variants in the 1 million genome era.

## Acknowledgements

Dr Andrea Talenti is funded by the Jamieson Bequest fund at the School of Biodiversity, One-Health and Veterinary Medicine of the University of Glasgow. The work has been made possible by the MARS High-Performance Compute cluster of the College of Medical, Veterinary and Life Sciences of the University of Glasgow.

## Captions

Supplementary Figure 1: Comparison of the Manhattan plots generated from a) XPCLR, b) XPCLRS and c) XPCLRS in `--fast` mode. The plots show the equivalency of the results produced from XPCLRS, and the more marked differences introduced by the non-adaptive integration in fast mode.

Supplementary Figure 2: Comparison of XPCLR and XPCLRS results when using PLINK inputs. The results are only unphased, and with/without the --fast option, showing the consistency in the results achieved.

Supplementary Table 1: Table of the results from XPCLR and XPCLRS for the AFR vs EUR comparison.

Supplementary Table 2: Performance comparison of the analyses ran by XPCLR and XPCLRS with the different configurations.

