## Supplementary figures and images for "XPCLRS: fast selection signature detection using cross-population composite likelihood ratio"

### Supplementary Figure 1

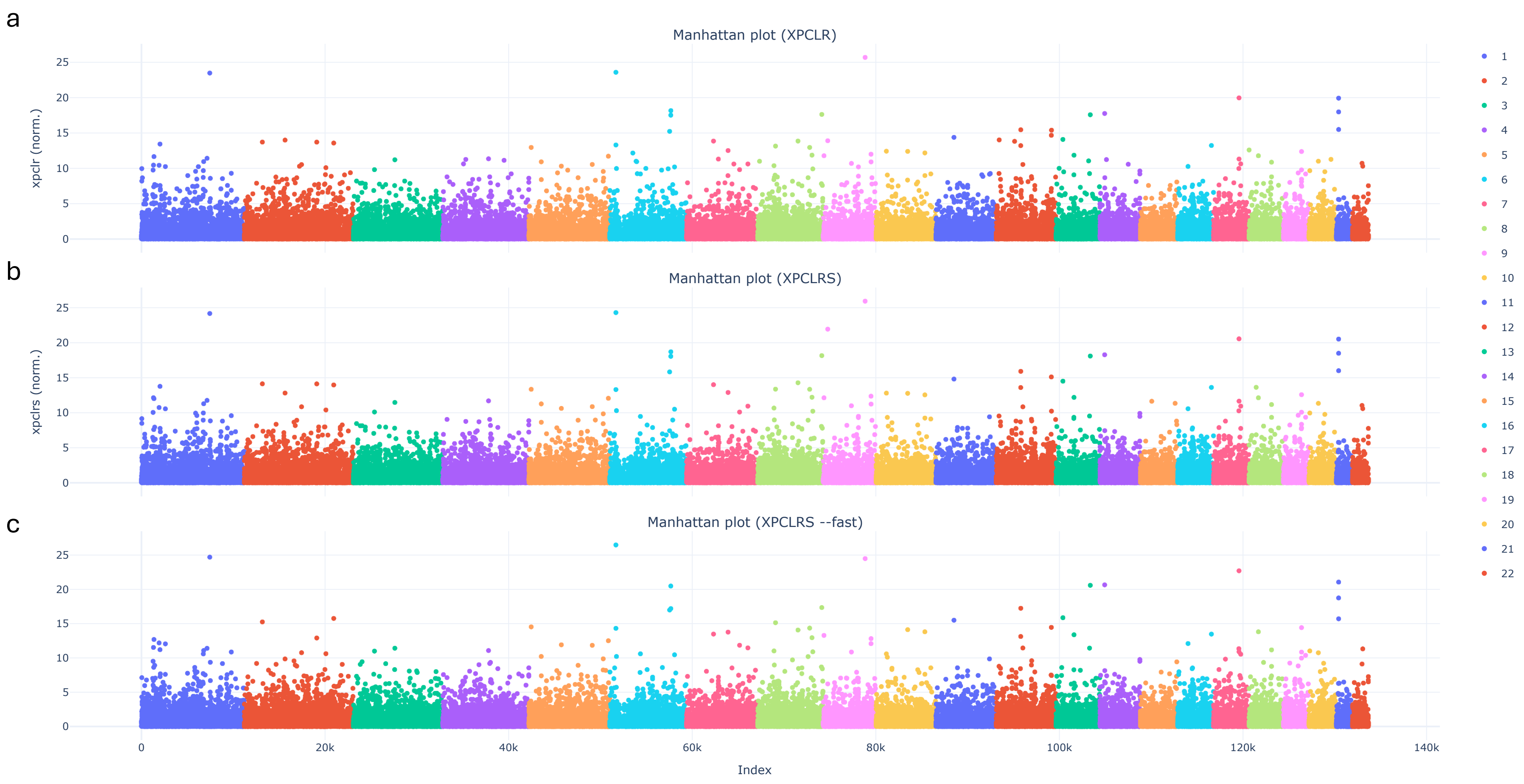

### Supplementary Figure 2

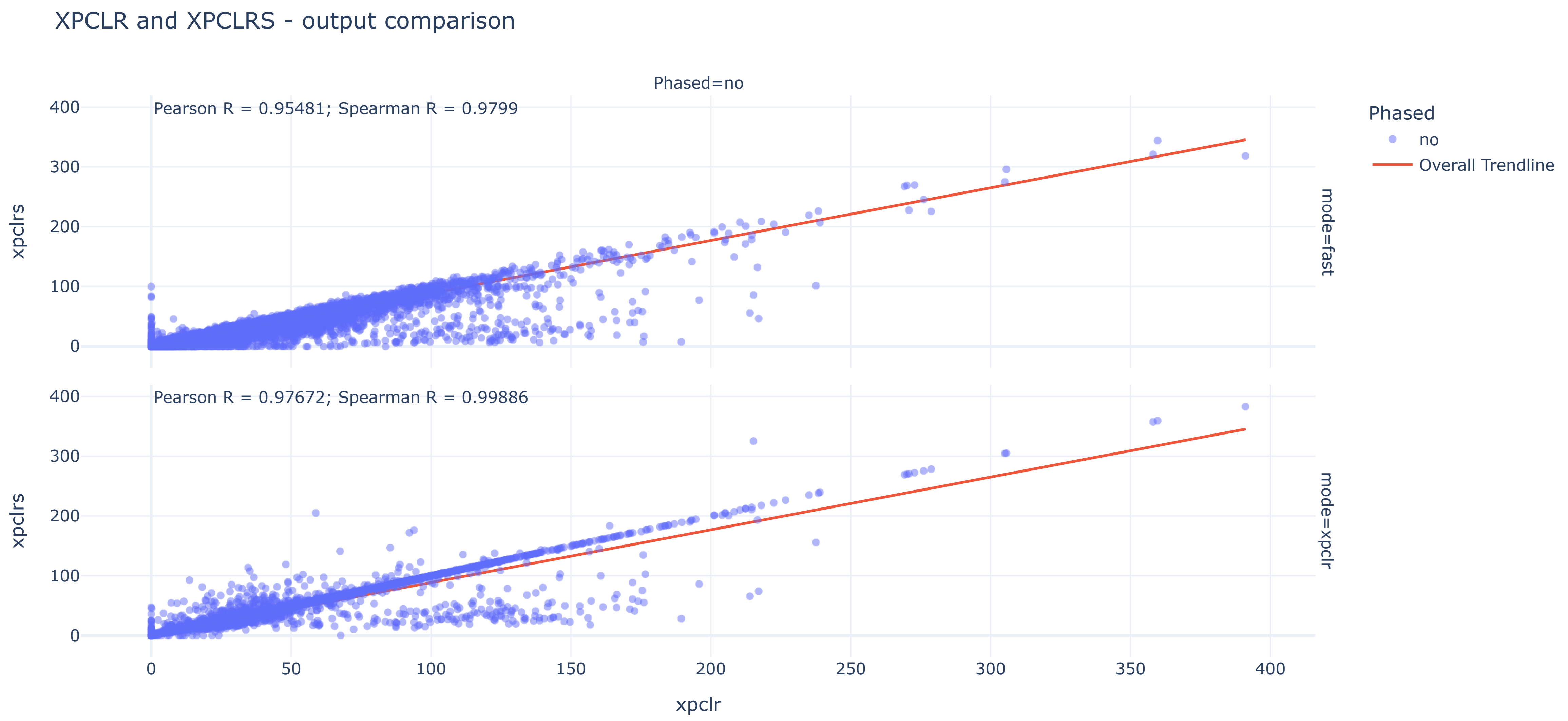
